# MORPhA Scale: behavioral and electroencephalographic validation of a rodent anesthesia scale

**DOI:** 10.1101/475921

**Authors:** Madalena Esteves, António M Almeida, Joana Silva, Pedro Silva Moreira, Emanuel Carvalho, José Miguel Pêgo, Armando Almeida, Ioannis Sotiropoulos, Nuno Sousa, Hugo Leite-Almeida

## Abstract

Degree of anesthesia in laboratory rodents is normally evaluated by testing loss of reflexes. While these are useful endpoint assessments, they are of limited application to study induction/reversal kinetics or factors affecting individual susceptibility (e.g. sex or age). We developed and validated a grading system for a temporal follow up of anesthesia. The Minho Objective Rodent Phenotypical Anesthesia (MORPhA) scale was tested in mice and rats anaesthetized with a mixture of ketamine/dexmedetomidine (ket/dex - 3 doses). The scale comprises 12 behavioral readouts organized in 5 stages – (i) normal/(ii) hindered voluntary movement, elicited response to (iii) non-noxious/(iv) noxious stimuli and (v) absence of response – evaluated at regular time points. Progression across stages was monitored by electroencephalography (EEG) in rats during anesthesia induction and reversal (atipamezole) and during induction with a second anesthetic drug (pentobarbital). Higher anesthetic doses decreased the time to reach higher levels of anesthesia during progression, while increasing the time to regain waking behavior during reversal, in mice and in rats. A regular decrease in high frequencies (low and high gamma) power was observed as the MORPhA score increased during anesthesia induction, while the opposite pattern was observed during emergence from anesthesia through reversion of dex effect. The devised anesthetic scale is of simple application and provides a semi-quantifiable readout of anesthesia induction/reversal.

## 1 Introduction

Anesthesia is routinely used in a number of procedures using laboratory animals. These studies, mostly in rodents, have been instrumental to understand the effectiveness of anesthetic drugs (Erhardt et al., 1984;Wixson et al., 1987;Hu et al., 1992;Field et al., 1993;Gardner et al., 1995), their biological mechanisms (Saunders and Ho, 1990;Accorsi-Mendonca et al., 2007) and potential adverse effects (Folle and Levesque, 1976;Erhardt et al., 1984;Field and Lang, 1988;MacDonald et al., 1989;Hu et al., 1992;Field et al., 1993;Janssen et al., 2004). Guidelines for rodent anesthesia are commonly available (Gaertner et al., 2008;Hellebrekers and Hedenqvist, 2010) and anesthetic regimen varies according to the species, strain and purpose being most commonly administered via injection or inhalation. Inhalational anesthesia (isoflurane, sevoflurane, halothane) allows controlled administration of drugs and fast reversal, though it requires dedicated equipment and specific expertise and safety conditions. On the other hand, injectable anesthetics are less expensive, requiring no equipment and are generally preferred. Commonly, they rely on associations of different drugs, allowing complementary effects at reduced dosages. For example, ketamine’s (ket) dissociative action can be complemented with a sedative/muscle relaxant such as xylazine or dexmedetomidine (dex). Nonetheless, for some purposes (e.g. electrophysiology in anesthetized animals) administration of a single drug such as pentobarbital may be preferable (Gaertner et al., 2008).

Proper monitoring of anesthesia conditions is of the highest importance to further explore the mechanisms that lead to effective anesthesia (Irifune et al., 2000;Solt et al., 2014;Zhou et al., 2015;Taylor et al., 2016), as well as to prevent unnecessary pain, discomfort and distress (Gaertner et al., 2008;Smith and Danneman, 2008;Hellebrekers and Hedenqvist, 2010), side effects known to impact on cognitive (Shields et al., 2016) and molecular read-outs (Black, 2002). In rodents, the presence/absence of reflexes to noxious stimuli (e.g. tail pinch) are used as a marker of proper anesthetic conditions (Smith and Danneman, 2008). While it provides a qualitative measure, it is not an effective monitoring procedure and lacks validity and reliability being a gross estimate and not a fine-tuned procedure. Alternatively, blood pressure and heart rate evaluation can be considered as fine markers of animal reactivity (sympathetic response to stress) (Smith and Danneman, 2008); these require specific equipment and might be difficult or even impractical in small animals and in certain types of surgeries/interventions (Smith and Danneman, 2008). Electroencephalography (EEG) monitoring, provides the most direct readout of anesthesia effect (Marchant et al., 2014). Indeed, electrophysiological correlates of commonly used behavioral outcomes such as righting reflex (MacIver and Bland, 2014;Pal et al., 2015) or paw pinch (Kortelainen et al., 2012) have been described for some drugs. However, these are normally used as anesthetized/awake endpoints and are not able to accompany the evolution of the anesthesia process. Moreover, currently described time (Bol et al., 1997;Vijn and Sneyd, 1998;Haberham et al., 1999;Hunt et al., 2006;Jang et al., 2009) and dosage-related (Bol et al., 1997;Hunt et al., 2006;Ihmsen et al., 2008;Jang et al., 2009) cortical dynamics do not fully elucidate the association between state of awareness and cortical activity. Additionally, EEG recordings in rodents require equipment, technical expertise and an additional surgery for electrode implantation which hinder its use for common procedures.

To fill in this gap we have designed the Minho Objective Rodent Phenotypical Anesthesia (MORPhA) scale, which comprises 12 behavioral readouts of anesthesia progression structured in 5 stages: (i) normal voluntary movement, (ii) abnormal voluntary movement, (iii) response to non-noxious stimuli, (iv) response to noxious stimuli and (v) absence of response to noxious stimuli. We evaluated MORPhA’s ability to discriminate different dosages of anesthetic (ket/dex) for induction and reversal (after atipamezole administration) of anesthesia in Wistar rats. Biological significance of the scores was assessed by comparison with EEG readouts. In order to further validate the scale, it was also used to evaluate ket/dex anesthesia and reversal in another commonly used laboratory animal, the B57BL/6J mouse. Additionally, rat MORPhA/EEG associations were established for pentobarbital, an anesthetic drug with a different mechanism of action.

## 2 Material and Methods

### 2.1 Animals

Six male Wistar-Han rats (Charles-River Laboratories, Barcelona, Spain) weighing 400-600 g (single-housed) and 5 male C57BL/6J mice 30-35 g were maintained at 22ºC temperature, 12 h light/dark cycle (lights on at 8 a.m.) and ad libitum food and water. All experiments were performed during the light phase of the cycle and conducted according to European Union Directives (2010/63/EU) and were approved by the institutional ethics commission - Subcomissão de Ética para as Ciências da Vida e da Saúde (SECVS).

### 2.2 EEG electrode placement

Animals (rats) were deeply anesthetized using a 2-4% sevoflurane in 100% oxygen mixture and placed in a stereotaxical frame where anesthesia and temperature were maintained through a nose mask and a heating pad, respectively. Stainless steel screws connected to golden Mill-Max receptacles (Mill-Max Mfg. Corp., Oyster Bay, NY, USA) were inserted in the skull in contact with the cortex in frontal (anterior-posterior (AP) +1.5, medial-lateral (ML) −2.0 relative to bregma), parietal (AP −1.0, ML +2.5), hippocampal (AP −4.3, ML −2.0) and occipital (AP −7.0, ML +3.0) areas, while two others in the cerebellum served as ground (AP −10.0 ML −2.0) and reference (AP −10.0 ML +2.0). These electrodes were connected to a Mill-Max connector and cemented to the skull. Animals were allowed to recover for one week before initiating experimental procedures.

### 2.3 Experimental design and anesthetics

Rats: 4 post-surgery experimental time-points were established at weekly intervals. On the first 3 weeks, 3 different dosages of ketamine/dexmedetomidine (ket/dex - Imalgene 1000®, Merial Portuguesa; Dorbene®, Zoetis Portugal) mix were intraperitoneally (i.p.) administered (60/0.340, 45/0.255 and 30/0.170 mg/Kg) in a latin square scheme (i.e. all animals received all three dosages at different time points). Application of the anesthesia scale started immediately after injection and was performed for 20 min, after which a unique dosage of the antagonist atipamezole hydrocloride (Antisedan®, Esteve Farma; 0.35 mg; i.p.) was injected and the scale was applied in the reverse order for 20 min. The responsible researcher was blind to the ket/dex dose. On the last week, a single dosage of sodium pentobarbital (Eutasil®, Ceva Saúde Animal) was administered (200 mg/Kg; i.p.) and the scale was applied for 20 min after which a lethal dosage of the same drug was administered. EEG was acquired in all trials (see below); 1 animal damaged the electrodes before the terminal experiment with pentobarbital and therefore EEG was not obtained. Temperature was not controlled due to potential interference with electrophysiological readouts and because the risk of hypothermia was low as experimentation time was brief. After sacrifice, brains were removed and visually inspected for surgery or infection-related damage.

Mice: 3 experimental time-points were established at weekly intervals, at which 3 different ket/dex dosages were administered (75/0.850, 50/0.565 and 25/0.281 mg/Kg; i.p.). The experimental protocol was applied as described for rats and 20 min after administration of the anesthetic mix, a fixed dose of atipamezole hydrochloride was injected (0.033 mg; i.p.). After locomotion was lost and until the end of the experiment, animals were kept in a heated environment.

### 2.4 Minho Objective Rodent Phenotypical Anesthesia (MORPhA) scale

The Minho Objective Rodent Phenotypical Anesthesia (MORPhA) scale construct created empirically but also based on procedures commonly applied in human loss of consciousness (Starmark et al., 1988). It is based on the observation of objective behavioral changes that represent progressive alterations of central nervous system function and corresponding reversion during recovery or the use of anesthetic antagonists. The MORPhA scale (Fig.1) was applied in rats and mice, starting at the moment of injection and for a total of 20 min. It consists of 12 score levels (from 0 to 11) divided in 5 stages: normal voluntary behavior (score 0), hindered voluntary movement (scores 1-3), elicited response to non-noxious stimuli (scores 4-6), elicited response to noxious stimuli (scores 7-10) and absence of response (score 11). For every evaluation a positive response (see Fig.1) led to the attribution of that score and re-assessment of the same behavior at the next indicated time-point, while absence of response entailed immediate evaluation of the behavior of the next score level.

**Fig. 1.**
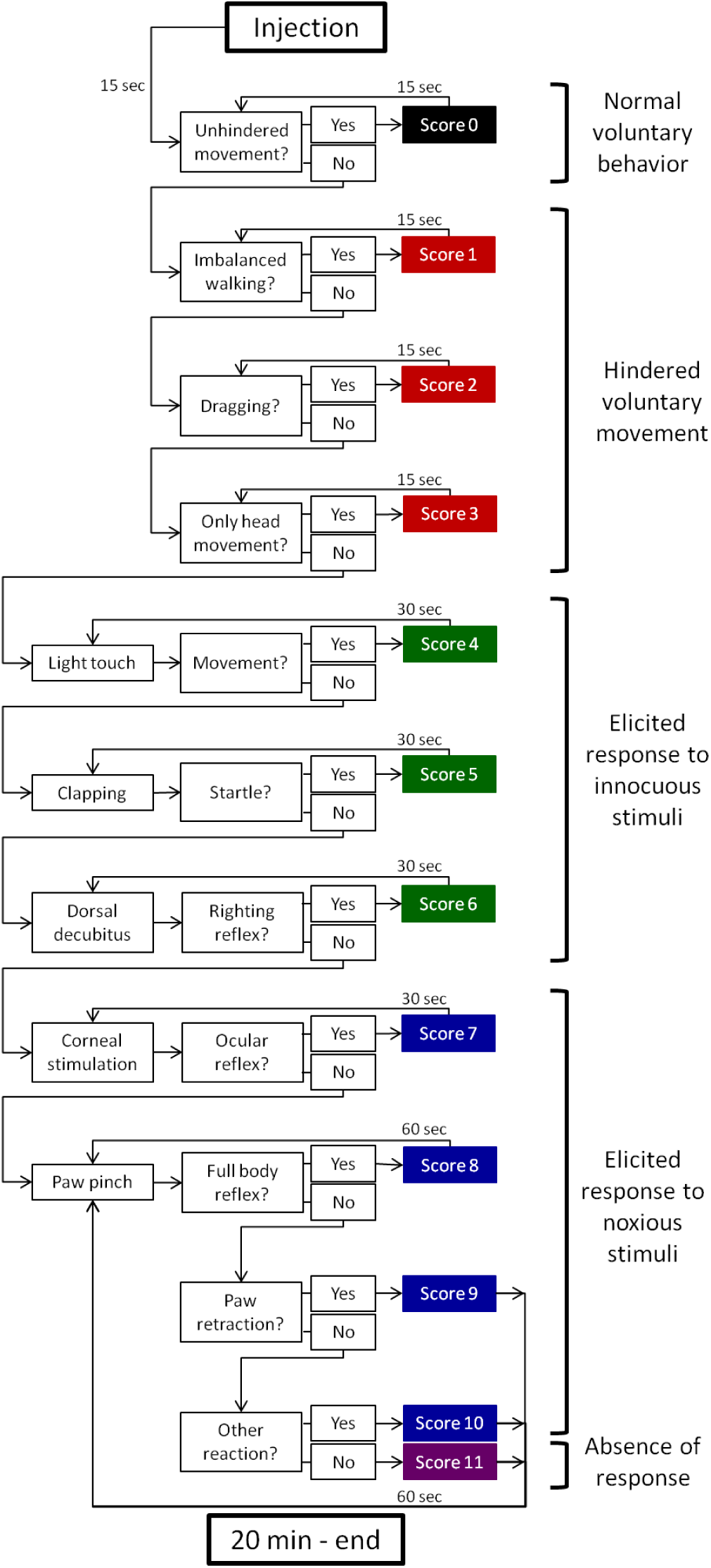
MORPhA scale. The scale is divided into 12 scores grouped in 5 stages – (i) normal voluntary behavior; (ii) hindered voluntary movement; (iii) elicited response to innocuous stimuli; (iv) elicited response to noxious stimuli; (v) absence of response. Animals are evaluated at 15 seconds intervals at the initial stages, where voluntary movement can be observed and thereon evaluations are spaced to 30 and 60 seconds with the increasing intensity of the stimuli. The MORPhA scale can be applied in both directions, i.e. for anesthesia induction and reversal.

Score 0 corresponds to normal, unhindered behavior and at this point reassessment was performed every 15 sec. Score 1 was attributed when the animal presented abnormal gait, i.e. it was able to ambulate in the cage using all four limbs but its movement was unbalanced. Score 2 required inability to lift the ventral region from the ground and consequent dragging of the body, while score 3 was reached when ambulation was not possible and only head movements were present. After no voluntary behavior was observed, stimulation was initiated and score 4 was attributed if any movement was observed after light touch by experimenter hand. At this point re-evaluations were executed at 30 sec intervals. Score 5 corresponds to a startle response after a loud hand clap over the animal’s head and score 6 to an attempt to recover position after placement in a dorsal decubitus position (righting reflex). If the animal did not respond to this last stimulus, noxious stimuli started was applied. Score 7 was attributed when ocular reflex was elicited after stimulation of the cornea with a cotton swab. The last stimulus of the scale was the paw pinch; it was administered with a bulldog forceps (jaw length 23 mm; AESCULAP), at 60 sec intervals in alternating posterior paws. Scores 8, 9 and 10 were ascribed after reactions to this stimulus, respectively full body reaction similar to a startle, paw withdrawal or subtle non-specific reaction such as alteration in breathing pattern. Score 11 corresponded to a total absence of response to this last stimulus.

### 2.5 EEG acquisition and analysis

On the day before the first experiment, rats were habituated to the recording site. The headstage was attached to the Mill-Max connector and animals were allowed to freely explore for 30 min. On recording days, animals were habituated for 5+5 min (headstage free/headstage connected) before each recording. A 5 min baseline was then allowed after which the anesthesia protocol was started. EEG data was acquired with the animal in its home cage using the dacqUSB system (Axona Ltd., London, UK) at 4800 Hz. Signals were amplified 3000 times and low-pass filtered at 150 Hz. All channels were referenced at the source and all EEG experiments were performed inside a Faraday cage.

Electrophysiological data was imported to Matlab (Mathworks, Natick,MA, USA) and analyzed with custom-written scripts using Chronux toolbox (http://chronux.org/) (Mitra and Bokil, 2008). Data was downsampled to 300 Hz and temporal correspondence between electrophysiological and behavioral recordings was established. Trials were defined as 7 sec before and after each score attribution, guaranteeing no overlap between trials. Trials in which the signal saturated were removed. Posterior visual inspection of the spectrum revealed interferences at 25 and 75 Hz which were associated with manipulation of the animals. Because removal in the time domain was not possible, 22-28 and 72-78 Hz frequencies were removed from further analysis.

Power estimates for each trial were attained by calculating the area under the curve (AUC) of the spectral density estimated with a multitaper method using Chronux (window 1.5 sec with 50% overlap, time-bandwidth product (TW) 3, number of tapers (K) 5, with a final frequency resolution of 0.59 Hz). Representation of each stage of anesthesia (scores 0, 1-3, 4-6, 7-10 and 11) for each animal was calculated as the mean AUC of every trial of that stage normalized for animals’ baselines (50-250 sec after the beginning of the recording, i.e. before administration of the anesthetic). Average power spectra of all animals for anesthesia with ket/dex, reversal and pentobarbital anesthesia is represented in Figs. 3-5A.

**Fig. 3.**
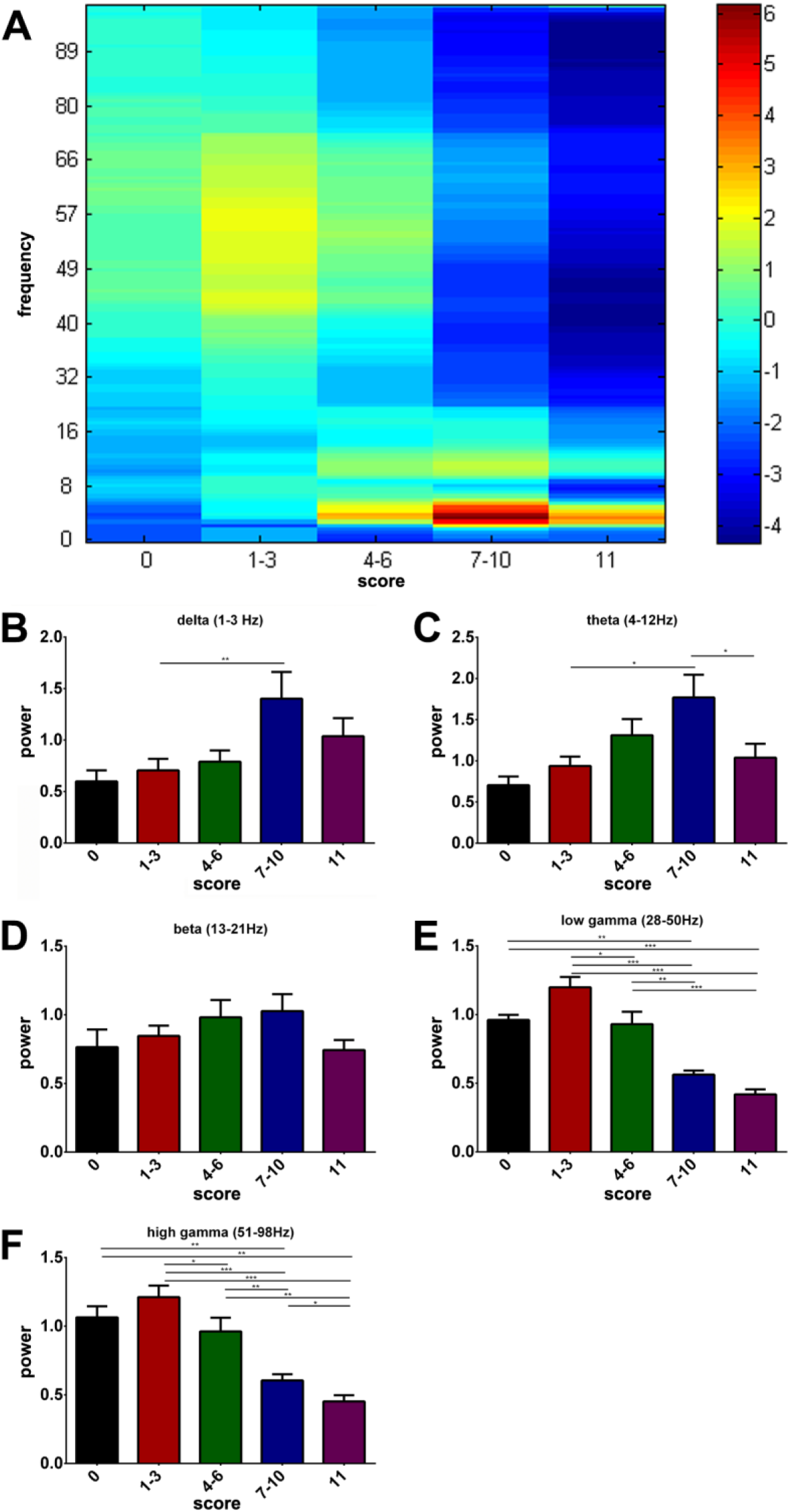
EEG activity across scale’s stages in anesthesia induction with ketamine/dexmedetomidine. (A) General power estimate spectrum showing power from 0 to 98 Hz for each of the five stages of anesthesia. (B-D) Average power for each anesthesia stage within five ranges of frequency: delta (1-3 Hz – B), theta (4-12 Hz – C), beta (13-21 Hz – D), low gamma (28-50 Hz – E) and high gamma (51-98 Hz – F). Statistics shown as asterisks corresponds to the analysis shown in Table 1, i.e. intra-animal pairwise comparisons. *p<0.05, **p<0.01, ***p<0.001 after Bonferroni correction for multiple comparisons.

For within frequencies analyses, ranges were defined as: delta 1-3 Hz, theta 4-12 Hz, beta 13-21 Hz, low gamma 28-50 Hz and high gamma 51-98 Hz (excluding 72-78 Hz).

**Table 1.**
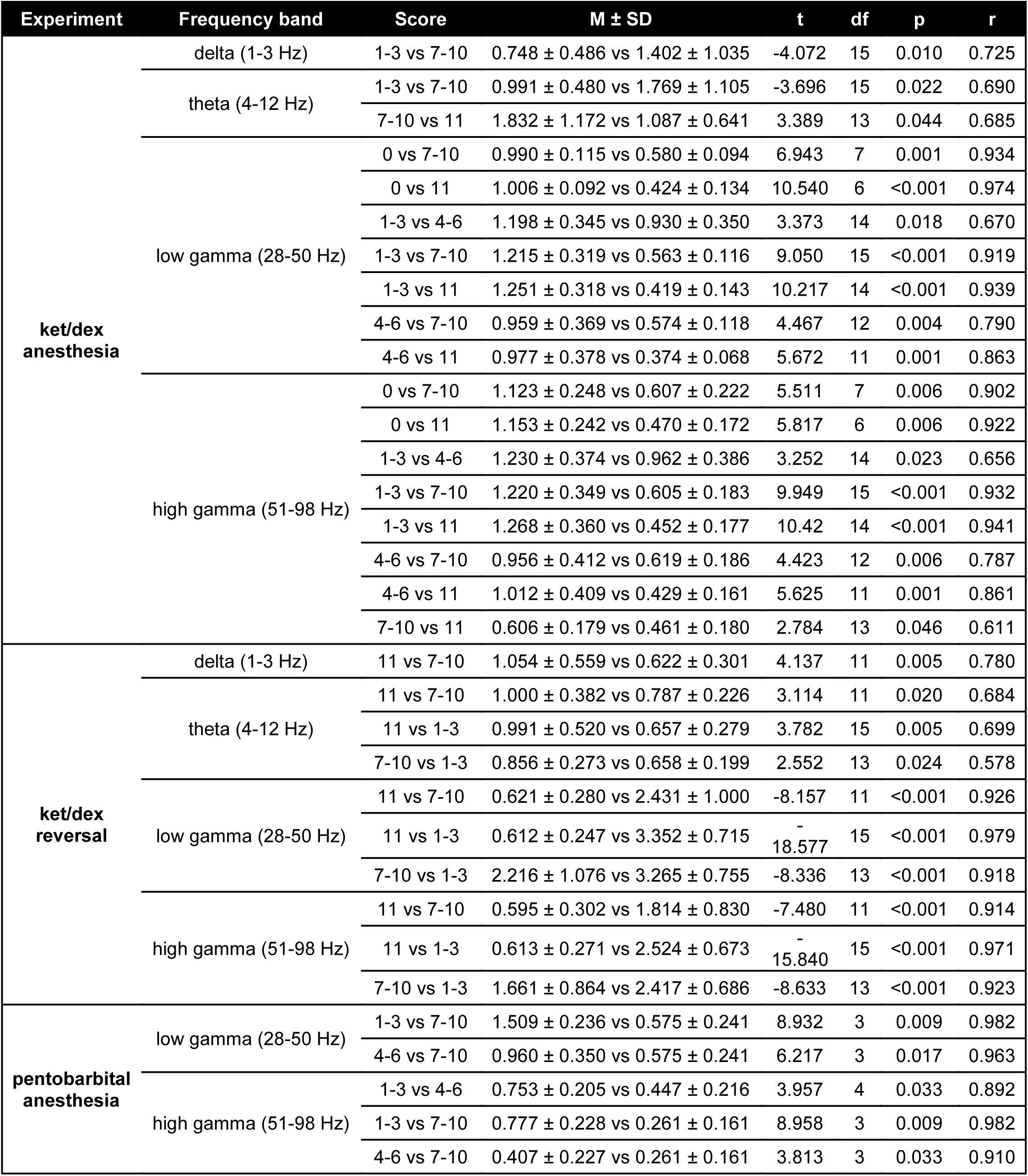
Statistically significant results of anaesthesia stages’ pairwise comparisons for ketamine/dexmedetomidine anaesthesia, respective reversal with atipamezole and pentobarbital anesthesia. M – mean; SD – standard deviation; t – student’s t; df – degrees of freedom; p – Bonferroni corrected p-value; r – effect size

### 2.6 Statistical analysis

All statistical analyses were performed in Matlab2009b software (The MathWorks, Inc., Natick, Massachusetts, United States). Power estimate spectra were attained in the same software and all remaining graphs were computed in Prism6 (GraphPad Software, Inc., La Jolla, USA). Mean and SEM of all animals included in the within frequencies analysis are shown in the bar graphs (Figs. 3-5B-F). As not all animals went through all scores (particularly in the highest doses where progression was fast), EEG analysis was divided into 5 stages of anesthesia and intra-animal pair-wise comparisons were performed for each range of frequencies (datasets comprehending n<4 were excluded from statistical analyses). As electrophysiological patterns were similar in all channels, only the occipital readings are shown. Data related to other channels can be found in supplementary Fig. S1. This statistical analysis is shown in the asterisks of the bar graphs and on the Table (Figs. 3-5B-F, Table 1). A power estimate spectrum for each of these comparisons was also extracted (see supplementary Fig. S2). Normality was verified using the Shapiro-Wilk test and paired t-tests were used for comparison between stages of anesthesia within each frequency range. Bonferroni-Holm correction for multiple comparisons was performed in all analyses and results were considered significant if corrected p-value was <0.05. Anesthesia progression and reversal curves are represented as mean and standard error of the mean (SEM) score of the previous minute. Mixed ANOVA with dose as between- and time as within-subjects factors was performed in order to evaluate dose-dependent differences in the curves.

## 3. Results

### 3.1 MORPhA scale is able to accompany anesthesia induction and reversal in both mice and rats

Administration of ket/dex induced a time- and dose-dependent progression in MORPhA score in rats (dose effect: F(2,15)=2.137, p=0.153; dose/time effect: F(36,270)=2.011, p=0.001). The top dosage of ket/dex (60/0.34 mg/Kg), led to a rapid progression to score 11 (absence of evoked reflex to a strong noxious stimulus) in all animals, while intermediate and low dosages (45/0.26 and 30/0.17 mg/Kg) induced an expected slower progression with some animals (1 out of 6 for each dosage) failing to reach score 11 (Fig. 2A). Administration of atipamezole induced a reversal of the MORPhA score (Fig. 2C) which was faster in animals treated with lower anesthetic dosages (dose effect: F(2,15)=2.907, p=0.086; dose/time effect: F(36,270)=1.524, p=0.034). During the time of evaluation, the minimal observed score was 1 (no animal recovered to score 0).

**Fig. 2.**
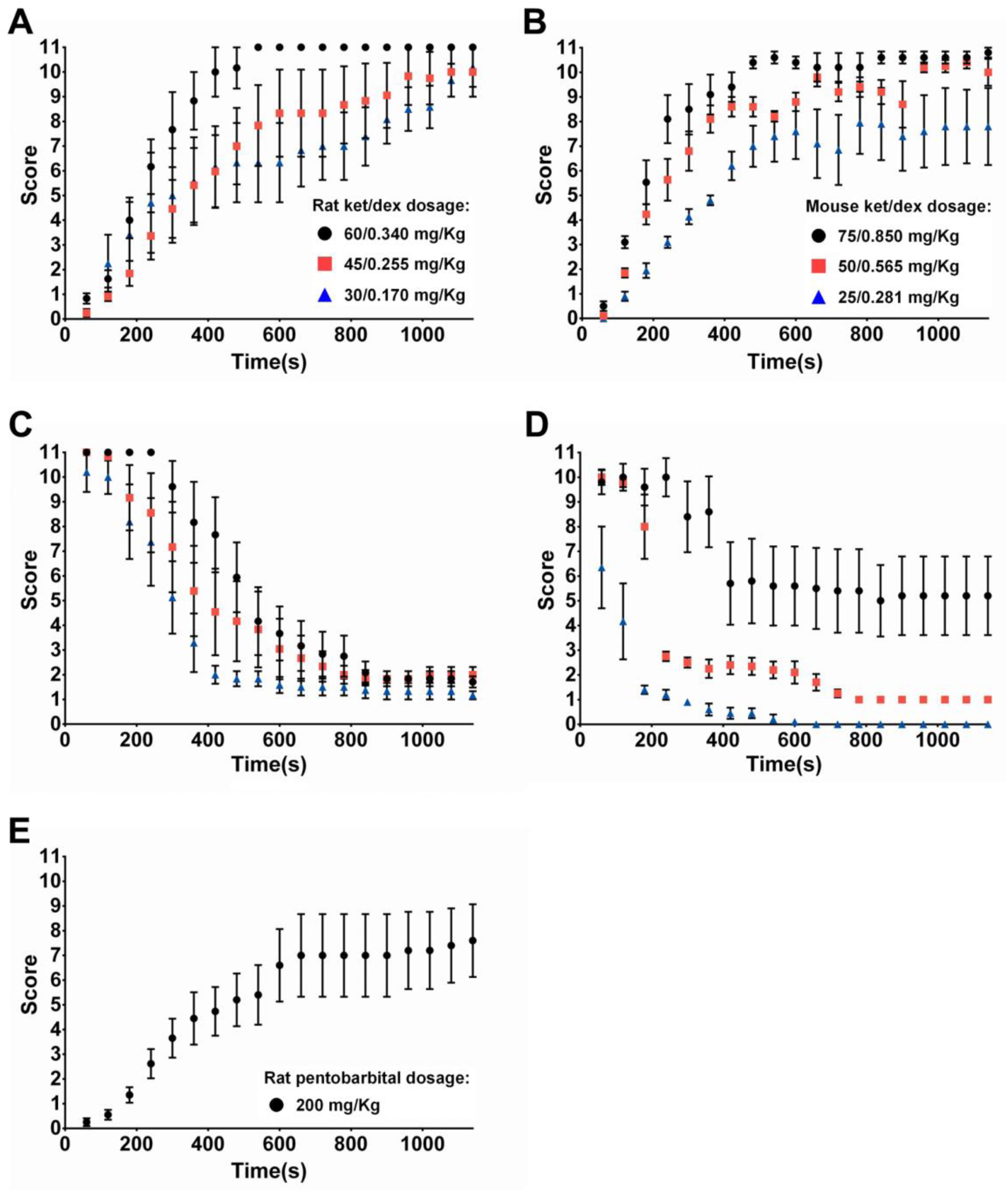
MORPhA progression during anesthesia induction and reversal. (A) Increasing doses of ketamine/dexmedetomidine (30/0.17, 45/0.26 and 60/0.34 mg/Kg) produced a leftward shift in anesthesia induction curve in rats. Conversely, respective (C) reversal with atipamezole hydrochloride (0.35 mg) was slower. (E) Pentobarbital administration (200 mg/Kg) also induced a progressive increase in MORPhA score, although most animals did not reach score 11. (B) As in rats, increasing doses of ketamine/dexmedetomidine (25/0.281, 50/0.565 and 75/0.850 mg/Kg) produced a leftward shift in anesthesia induction curve in mice. Conversely, respective (D) reversal with atipamezole hydrochloride (0.033mg) was slower.

Also, in mice, the MORPhA scale was able to accompany progression of anesthesia and to effectively distinguish the 3 ket/dex dosages (75/0.850, 50/0.565 and 25/0.281 mg/Kg) (Fig.2>B - dose effect: F(2,12)=10.275, p=0.003; dose/time effect: F(36,216)=0.964, p=0.533). Moreover, administration of a fixed dose of antagonist (atipamezole) induced anesthesia reversal that was dependent on ket/dex dosage (Fig.2D - dose effect: F(2,12)=15.538, p<0.001; dose/time effect: F(36,216)=4.225, p<0.001). At the end of the experiment, and taking advantage of the time of sacrifice, a sub-lethal dosage of pentobarbital was administered. Although on average the animals did not reach score 11, the MORPhA scale was also able to accompany this progression (Fig. 2E).

### 3.3 MORPhA ket/dex-induced progression is associated with EEG power in rats

The occipital channel will be used to illustrate scale correlates – frontal, parietal, hippocampal and occipital readings provided similar results (consult supplementary Fig.S1 for the complete data set). Visual inspection of the general power estimate spectrum (Fig. 3A) shows a pattern that can be analyzed within ranges of frequencies (Fig. 3B-F, Table 1). The overall view is a progression of increasing power in the delta (1-3Hz), theta (4-12Hz) and beta (13-21Hz) range and a decrease in power in the gamma (low 28-50Hz; high 51-98Hz) frequencies as is depicted in the power estimate spectra. Power in the delta range (Fig. 3B) significantly increased in scores 7-10 when compared with scores 1-3 (Table 1). In theta frequencies (Fig. 3C), power gradually increased until scores 7-10 and finally decreased when the animal was completely non-respondent (Table 1). A similar although not statistically significant pattern was found for the beta range (Fig. 3D). In both low and high gamma frequencies, there was an obvious power-score association and a gradual decrease in power until score 11 was reached could be observed (Fig. 3E-F, Table 1). Consult supplementary Fig. S2 for power estimate spectra of significant comparisons.

### 3.3 Anesthesia reversal-induced MORPhA score reduction is associated with EEG power in rats

As previously mentioned, none of the rats recovered to score 0 during the time of evaluation. Additionally, scores 4-6, which are shown in the general power estimate spectrum (Fig. 4A), were also excluded from the analysis due to their low representation. The pattern of evolution mostly mirrored the ones observed in the ket/dex induction phase but presented some differences, especially in the lower ranges of frequencies (Fig. 4). In fact, although the power for both delta and theta ranges for score 11 was very similarly symmetrical in both the induction and recovery phases (Figs. 4B-C and 3B-C), scores 7-10 and 1-3 suffered some alterations. In the delta range score 11 presented a higher power than scores 7-10 during reversal (Fig. 4B, Table 1), while during the induction phase scores 7-10 presented the highest power (Fig. 3B, Table 1). In the theta range, a gradual decrease in power was seen from score 11 until scores 1-3 (Fig. 4C, Table 1), which differs from the pattern observed during the anesthesia process (Fig. 3C, Table 1). Once again, in the beta range, no significant differences were found (Fig. 4D). In the gamma frequencies, power also presented a similar order of magnitude in score 11 when comparing with the induction phase (Figs. 4E-F and 3E-F) and reduction of score was associated with an increase in power (Fig. 4E-F, Table 1), although this increase was higher than seen in the previous experiment. Consult supplementary Fig. S2 for power estimate spectra of significant comparisons.

**Fig. 4.**
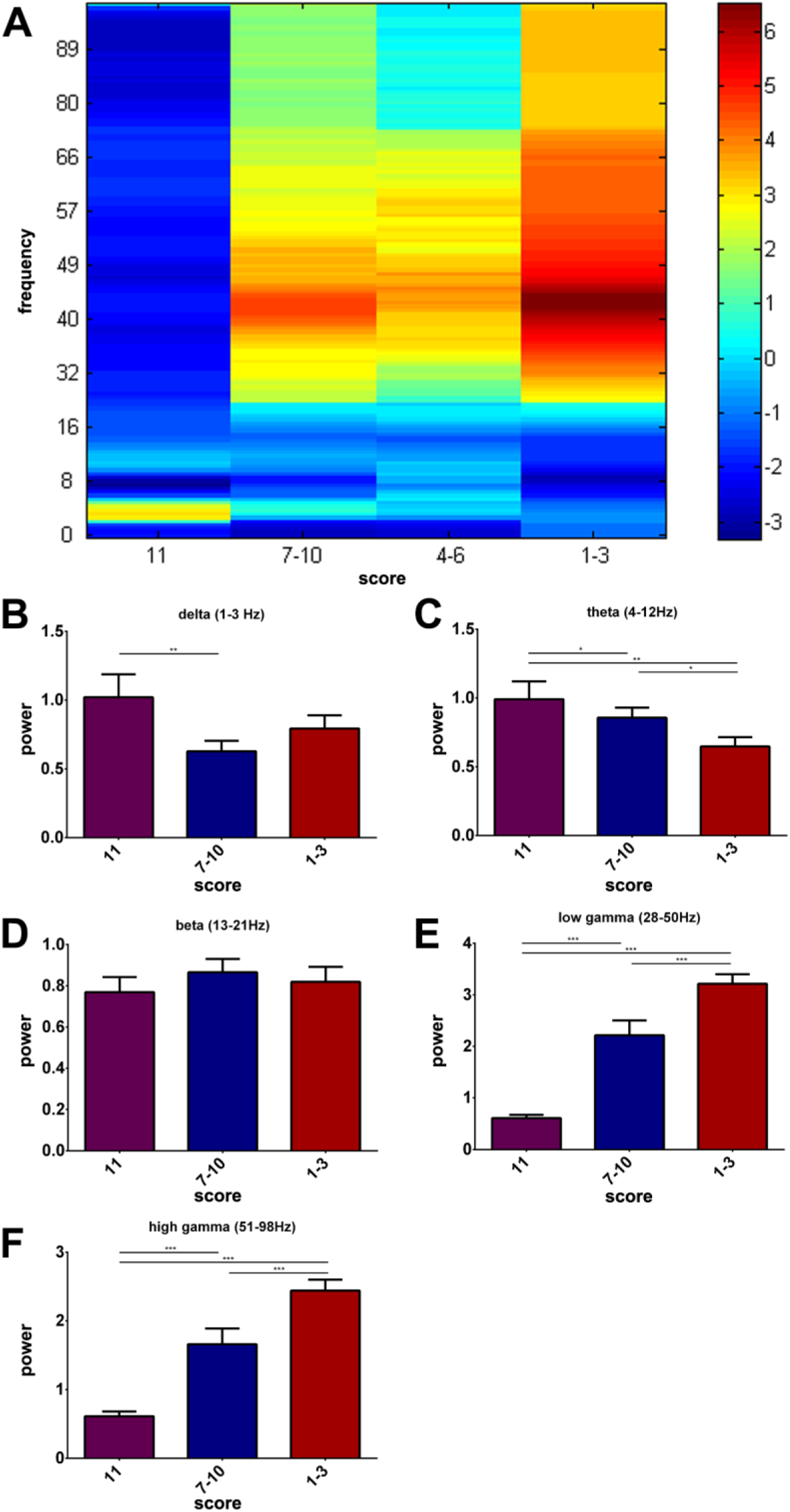
EEG activity across scale’s stages in anesthesia reversal with atipamezole. (A) General power estimate spectrum showing power from 0 to 98 Hz for four of the five stages of anesthesia (none of the animals reached score 0). (B-D) Average power for each of the analyzed anesthesia stage within five ranges of frequency: delta (1-3 Hz – B), theta (4-12 Hz – C), beta (13-21 Hz – D), low gamma (28-50 Hz – E) and high gamma (51-98 Hz – F). Statistics shown as asterisks corresponds to the analysis shown in Table 1, i.e. intra-animal pairwise comparisons. *p<0.05, **p<0.01, ***p<0.001 after Bonferroni correction for multiple comparisons.

### 3.4 MORPhA score correlates with EEG power induced by a different drug (pentobarbital)

Pentobarbital administration (200 mg/Kg i.p.) also induced a progression in the scale although there was high variability and most animals did not reach score 11. After drug administration, a general decrease in power can be observed in the power estimate spectrum (Fig. 5A). Due to low representation, comparisons with scores 0 and 11 were not possible, but comparisons between other scores showed no differences in the lower ranges of frequencies (delta, theta and beta; Fig. 5B-D). In low and high gamma ranges, a gradual decrease in power was observed as anesthesia progressed (Fig. 5E-F, Table 1). Consult supplementary Fig. S2 for power estimate spectra of significant comparisons.

**Fig. 5.**
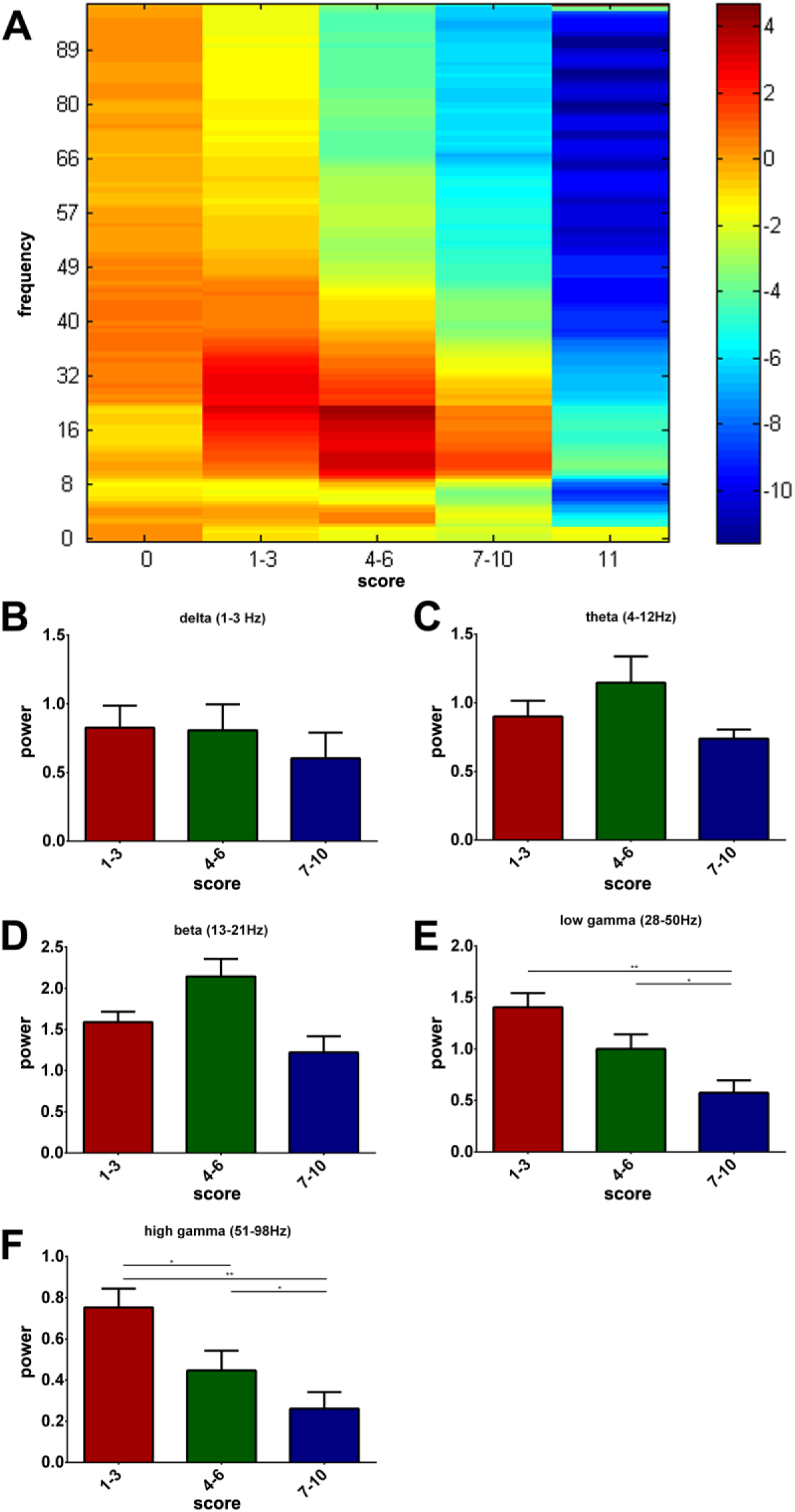
Effect of pentobarbital administration on MORPhA scale and correspondent EEG activity across the scale’s stages. (A) General power estimate spectrum showing power from 0 to 98 Hz for each of the five stages of anesthesia. (B-F) Average power for each of the analyzed anesthesia stages within five ranges of frequency: delta (1-3 Hz –B), theta (4-12 Hz – C), beta (13-21 Hz – D), low gamma (28-50 Hz – E) and high gamma (51-98 Hz – F). Statistics shown as asterisks corresponds to the analysis shown in Table 1, i.e. intra-animal pairwise comparisons. *p<0.05, **p<0.01, ***p<0.001 after Bonferroni correction for multiple comparisons.

## 4 Discussion

In this study we validated a new anesthesia scale in both rats and mice. The scale is of simple application in both species and provides a quantifiable measure of anesthesia induction and reversal. Additionally, in rats we also validated with EEG electrophysiological correlates of the anesthesia induction stages for two different drugs and reversal dex. We were able to find clear power/scale score associations, especially in higher frequencies, providing biological meaningfulness to the scale’s scores/stages and further validating the MORPhA scale.

The behavioral scale developed and validated for monitoring the anesthesia process in laboratory rodents is based on timely evaluations that retrieve scores from 0 to 11, where 0 corresponds to normal unhindered behavior and 11 corresponds to total absence of response to a strong noxious stimulus. Total experiment time was limited to 40 min, preventing observation of spontaneous reversal of anesthesia or complete reversal after atipamezole injection. It provided uniformization across experiments and allowed to perform the experiments in rats without the introduction of additional electronic devices for temperature control which could potentially interfere with the electrophysiological signal.

Besides the usage of anesthesia’s end-points such as presence/absence of response to tail pinch (Smith and Danneman, 2008), efforts towards building a scale that is able to determine anesthesia effectiveness have been put forward (Devor and Zalkind, 2001). However, MORPhA is the first to provide a systematic assessment with grading detail, in which not only is stimulus intensity adjusted to sedation levels, but also time interval of stimulus application is adjusted to its intensity, aiming to decrease the effect of scale application in normal anesthesia progression. As a result, this scale was able to accompany induction and reversal of anesthesia in both rats and mice and to distinguish different dosages of anesthetic mix (ket/dex) in all conditions. The moment of sacrifice was also used to assess the applicability of the scale to a different commonly used anesthetic - pentobarbital - by administering a non-lethal dosage. The MORPhA is therefore applicable in 2 anesthetic protocols commonly employed in laboratory environment and it has probably a wider range of applications. Nevertheless, its application in the context of other anesthetics, particularly gaseous, might require adaptation. Also, experimental designs involving very young/old animals or females subject to additional validation due to possible pharmacokinetic and/or pharmacodynamic specificities.

Electrophysiological validation was based on stages of anesthesia comprising biologically similar groups of scores: normal voluntary behavior, hindered voluntary movement, elicited response to innocuous stimuli, elicited response to noxious stimuli and absence of response. This classification was necessary due to the quick progression in the scale, particularly at higher dosages, which would not allow a good statistical power in the analysis. Nonetheless, each of these stages showed to be distinct in power from all others, further validating this division. No area-dependent differences were however found in the EEG profiles during anesthesia progression, which were similar in all recorded channels (frontal, parietal, occipital and hippocampal). Spatial progression has been reported in humans, showing a marked anteriorization of low frequency waves (John et al., 2001;John and Prichep, 2005) while high frequency oscillations increase in frontal areas and then migrate to posterior regions (John and Prichep, 2005). However, due to the small size of the animal’s brain and the relative large size of the implanted electrodes no differences were observed in the recorded channels. In fact, Jang and colleagues have previously reported similar results, with comparable recordings in frontal, parietal and occipital electrodes during rat ket/dex anesthesia (Jang et al., 2009).

Ket/dex mix injection induced an increase in power at lower frequencies (delta and theta) until scores 7-10 followed by a decrease on score 11. In contrast, higher frequencies (low and high gamma) power progressively decreased until score 11. During reversal lower frequencies showed a decrease in power that accompanied the awakening of the animal, while higher bands showed a gradual increase in power superior to baseline levels. Interestingly, the transition was often abrupt, passing from 11 to 2-3 scores in few seconds A previous study showed that ket/dex injection in anesthetic dosages induce an obvious (although not statistically evaluated) increase of power in frequencies under 22 Hz over time-points comparable with ours (0 to 20 min). This difference tended to be bigger at higher dex doses and to decrease with time (up to 70 min), while no obvious alterations were seen in the first 20 min at higher frequencies (22-50 Hz). These authors were also able to correlate response to paw pinch to power at 7-10 Hz and in accordance with our data, deeper anesthesia was associated with higher power (Jang et al., 2009). Reversal of this anesthesia induced an increase in power in frequencies between 14 and 50 Hz (vs baseline) (Jang et al., 2009), which is comparable to the increase we see between 25 and 100 Hz.

These differences between the process of anesthesia and reversal could eventually be attributed to the mechanism of action of atipamezole (Virtanen et al., 1989). This drug is able to revert the effects of dex (α2 agonist) but not of ket (methyl-D-aspartate antagonist) which could potentially explain the incomplete reversal of both behavioral (none of the rats reached score 0 in 20 min) and electrophysiological parameters. However, spontaneous recovery of ket-induced anesthetized state decreased power of the 0.5-4 Hz band and increased it in higher frequencies’ (up to 250 Hz) power, even when comparing to baseline levels. Initial electrophysiological state was only achieved after the animal recovered to normal behavior and pre-frontal acetylcholine levels returned to normal (Pal et al., 2015). Moreover, dex alone is also able to induce a dose-dependent increase in 0.5-3 Hz power whose reduction after plasma concentration decrease is delayed, i.e. similar concentrations during induction and reversal are associated with increased power in the latter (Bol et al., 1997). These “awakening” discrepancies between pre- and post-anesthesia should not therefore be associated with partial reversion of the process.

Before euthanizing the animals, a sub-lethal dosage of pentobarbital was administered, and scale/EEG correlates were also evaluated to maximize data collection. Previous reports have shown both a general increase in power up to 25 Hz (Haberham et al., 1999) during anesthesia with the same drug, as well as a decrease in 6-7 Hz power accompanied by an increase in the 2-3 Hz range (Devor and Zalkind, 2001). However, in our study, higher bands of frequency (25-100 Hz), which are normally overlooked, showed higher predictive value. As seen with ket/dex anesthesia, power in these bands decreased with the progression of anesthesia, while variations in lower frequency ranges could not be shown.

Different electrophysiological parameters have also shown associations with degree of anesthesia. In fact, administration of ket reduces overall coherence (Pal et al., 2015) while ket/dex mix induces a reduction in spectral edge frequency 90 and 95 (Jang et al., 2009). On the other hand, the effect of other anesthetics on brain waves has also been shown in the literature. Isoflurane anesthesia for example, has been shown to be strongly associated with a decrease in low (MacIver and Bland, 2014) and an increase in high frequency power (Kortelainen et al., 2012;MacIver and Bland, 2014), as well as with other parameters such as alterations in attractor shapes (MacIver and Bland, 2014), median (Ihmsen et al., 2008;Kortelainen et al., 2012) and spectral edge frequencies (Ihmsen et al., 2008;Kortelainen et al., 2012), spectral entropy (Kortelainen et al., 2012), relative beta ratio (Kortelainen et al., 2012) and approximate entropy (Ihmsen et al., 2008).

In summary, to the best of our knowledge, the electrophysiological correlates of anesthesia progression (and reversal) have never been assessed. In fact, the above-mentioned evaluations have been performed at specific cut off points of anesthesia and/or recovery while many others assess time- or dosage-dependent electrophysiological changes without behavioral correlates (see for instance (Ihmsen et al., 2008;Jang et al., 2009;Silva et al., 2010;Yoon et al., 2011;Flores et al., 2015)). Our scale was thus the first to show its ability to accompany this progression at both behavioral and electrophysiological levels. Despite potential necessity for adaptation to different anesthetic regimens, it was able to behaviorally distinguish different anesthetic dosages during anesthesia and reversal in both rats and mice and the rat EEG profile followed a stable evolution that accompanied the behavioral evaluation. It can thus be a new and valuable tool for the evaluation of anesthesia progression (induction/reversal) and of parameters affecting individual susceptibility (e.g. sex, age or genetic background), requiring minimal equipment or prolonged training.

## Supporting information

## 6 Author Contributions

M.E. and H.L.A: conception of the scale and design of the study; M.E., A.M.A., I.S., J.S. and E.C.: acquisition of data; M.E., P.S.M. and H.L.A.: data analysis; M.E., H.L.A., J.M.P., A.A., I.S. and N.S.: data analysis and interpretation. M.E. and H.L.A.: preparation of the manuscript’s initial version. All authors revised and approved the final version of the manuscript and agree to be accountable for all aspects of the work.

## List of Abbreviations

AP: anterior-posterior
AUC: anterior-posterior
dex: dexmedetomidine
EEG: electroencephalography
i.p.: intraperitoneal
K: number of tapers
ket: ketamine
ML: medial-lateral
MORPhA scale: Minho Objective Rodent Phenotypical Anesthesia scale
SECVS: Subcomissão de Ética para as Ciências da Vida e da Saúde
SEM: standard error of the mean
TW: time-bandwidth product

## 9 Acknowledgments

We would like to thank Dr. Luis Ricardo Jacinto for his technical assistance and valuable comments.

