## Supplementary material for "MORPhA Scale: behavioral and electroencephalographic validation of a rodent anesthesia scale"

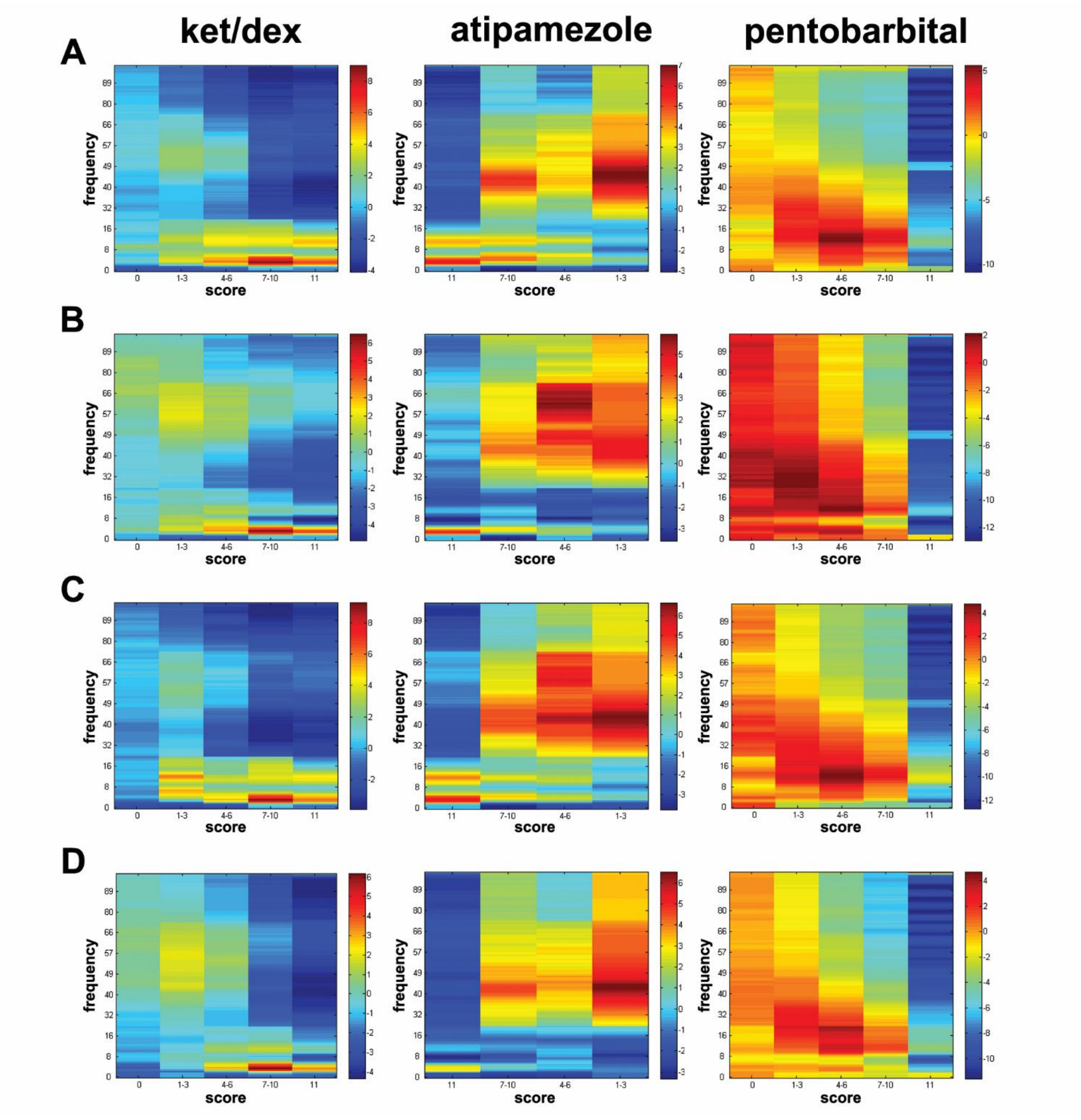


**Supplementary Fig. S1 General spectrograms of EEG activity in all recorded channels.** General spectrograms showing power from 0 to 98 Hz for each of the five stages of anesthesia after administration of ketamine/dexmedetomidine (ket/dex), atipamezole or pentobarbital in all recorded EEG channels - frontal, hippocampal, parietal and occipital.
