## Supplementary material for "MORPhA Scale: behavioral and electroencephalographic validation of a rodent anesthesia scale"

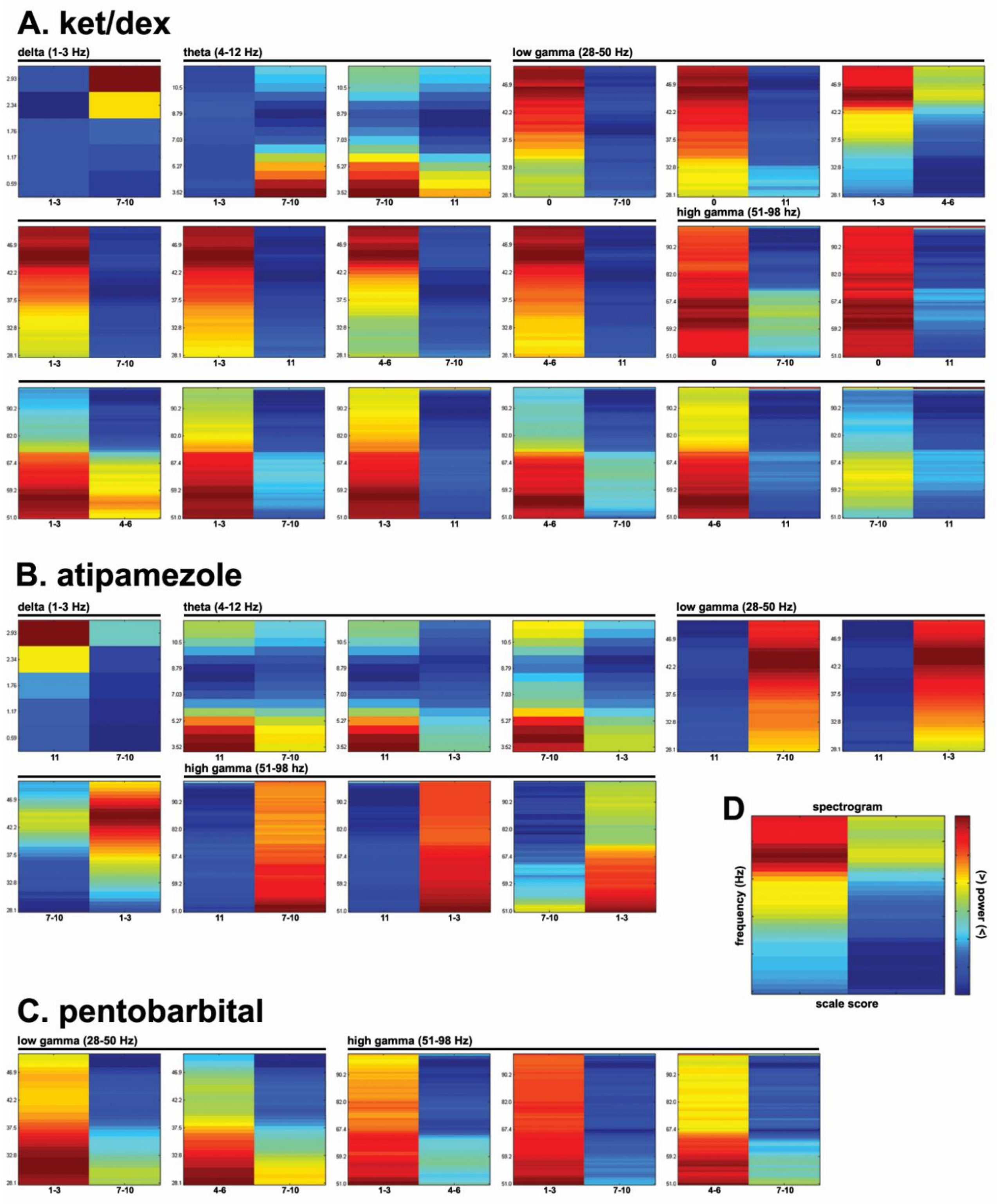


**Supplementary Fig. S2 Spectrograms of the significant intra-animal pair-wise comparisons.** For (A) ketamine/dexmedetomidine (ket/dex), (B) atipamezole and (C) pentobarbital administration, significant range-specific intra-animal comparisons of power between stages of anesthesia are represented. Representative legends are shown (D).
